# Spatial, climate, and ploidy factors drive genomic diversity and resilience in the widespread grass *Themeda triandra*

**DOI:** 10.1101/864298

**Authors:** CW Ahrens, EA James, AD Miller, NC Aitken, JO Borevitz, DJ Cantrill, PD Rymer

## Abstract

- Fragmented grassland ecosystems, and the species that shape them, are under immense pressure. Restoration and management strategies should include genetic diversity and adaptive capacity to improve success but these data are generally unavailable. Therefore, we use the foundational grass, *Themeda triandra*, to test how spatial, environmental, and ploidy factors shape patterns of genetic variation.
- We used reduced-representation genome sequencing on 487 samples from 52 locations to answer fundamental questions about how the distribution of genomic diversity and ploidy polymorphism supports adaptation to harsher climates. We explicitly quantified isolation-by-distance (IBD), isolation-by-environment (IBE), and predicted population genomic vulnerability in 2070.
- We found that a majority (54%) of the genomic variation could be attributed to IBD, while 22% of the genomic variation could be explained by four climate variables showing IBE. Results indicate that heterogeneous patterns of vulnerability across populations are due to genetic variation, multiple climate factors, and ploidy polymorphism, which lessened genomic vulnerability in the most susceptible populations.
- These results indicate that restoration and management of *T. triandra* should incorporate knowledge of genomic diversity and ploidy polymorphisms to increase the likelihood of population persistence and restoration success in areas that will become hotter and more arid.

## Introduction

Grasses (Poaceae) are one of the most ecologically important vascular plant groups, making up 25% of the world’s vegetation (Shantz, 1954). They provide key ecosystem services that underpin environmental health (i.e. habitat and food sources for native wildlife, nutrient cycling and carbon sequestration), and carry significant economic value as they include four of the five major crops in terms of global production (Raven & Thomas, 2010). Grasses are essential constituents of several vegetation communities including grasslands, grassy woodlands, and alpine regions. However, grasslands and grassy woodlands have historically been under immense pressure from rangeland and agricultural uses (Eldridge *et al.*, 2016; Hopkins & Holz 2006), leading to the fragmentation of natural populations and reductions in genetic diversity (Harrison *et al.*, 2015). Today, only about 4.6% of the billions of hectares of grassland ecosystems remain worldwide (IUCN 2016). In Australia, grassland systems are the most poorly conserved and degraded communities (Hobbs & Yates, 2000), and are likely to experience major negative long-term effects. Many regions of Australia that support grasslands are becoming warmer, drier and increasingly fire prone under climate change, highlighting the importance of preserving genetic diversity and evolutionary potential (Dunlop *et al.*, 2012). However, most research on genetic diversity in grass species has generally been undertaken on those of agricultural importance (Buckler *et al.*, 2001) such as wheat, corn, rice, and sorghum, or those that are being developed for biofuels such as switchgrass (*Panicum* – Casler *et al.*, 2007; Harrison *et al.*, 2015) and sugarcane (*Miscanthus* – Vermerris, 2008). While research on species such as switchgrass have provided valuable insights into natural patterns of genetic diversity, adaptation across gradients, and the role ploidy plays between these lines of enquiry (Morris *et al.*, 2011; Lowry *et al.*, 2014, 2019; Grabowski *et al.*, 2014), major gaps in knowledge for other ecologically important grasses persist and continue to inhibit effective conservation management.

Genetic diversity is maintained within a species by a combination of selective (such as range shifts and natural selection) and neutral processes (such as gene flow, mutation, and genetic drift) (Futuyma, 2013). However, grasses often have complex evolutionary histories (Stebbins, 1956) influenced by factors such as clonality (Fischer & Van Kleunen, 2002), polyploidy (Keeler & Bradshaw, 1998), intrageneric hybridization, genome size, and different physiologies such as photosynthetic mechanisms (e.g. C3 versus C4) (Edwards *et al.*, 2010). These complex and often lineage-specific life histories can complicate our ability to project findings across species, meaning that the species-specific data needed for practitioners to make informed management decisions is often lacking. Perhaps the lack of research on ecologically important grass species and their complex life histories are not mutually exclusive. Regardless, information about how genetic diversity is distributed across habitats and environmental gradients, often reflecting selection and local adaptation, can help inform management and restoration strategies (Hoffmann *et al.*, 2015). This is particularly pertinent given grassland communities are already showing signs of climate stress, and empirical data is urgently needed to support adaptive management strategies that prepare grasslands for new climate challenges by maximising evolutionary potential. In addition, research that focuses on genetic diversity across species ranges can help identify populations vulnerable to climate stress, allowing practitioners to prioritise management that safeguards populations at risk. For example, genomic signals of selection can be used to predict climate-driven population declines (Bay *et al.*, 2018). Specifically, ‘genomic vulnerability’ of individual populations, defined as the mismatch between current and predicted future genomic variation inferring population susceptibility to the loss of genetic diversity and/or maladaptation, can help identify populations most at risk. As our ability to integrate geospatial and genomic resources continues to grow, so will the ability of researchers to identify genomic vulnerability in ecologically important species, providing practitioners with improved management frameworks for mitigating climate change effects on ecosystems by preserving patterns of endemism and maximising adaptive potential.

Grasses often display ploidy differences among populations across their natural range. Indeed, polyploidy is common among vascular plants with c. 35% of species characterised as having a recent history of polyploidy (Wood *et al.*, 2009). For many species, associations between ploidy and local environmental conditions reflect adaptation, a pattern which has been studied extensively in crop plants (Alix *et al.*, 2017). Further, it has recently been shown that niche differentiation occurs faster in polyploids than diploid relatives (Baniaga *et al.*, 2019). While the causes of polyploidy are poorly understood (Soltis *et al.*, 2010), whole genome duplication events have been shown to coincide with historical climate change events (Cai *et al.*, 2019), and patterns of allopolyploidy have been linked to changes in environment (Wagner *et al.*, 2019). The effects of polyploidy are increasingly evident, with gene expression levels shown to vary from tissue to tissue in polyploids compared to their diploid counterparts (Adams *et al.*, 2003), and polyploid species often having significant fitness advantages (Petit & Thompson, 1997; Bretagnolle & Thompson, 2001; Ramsey, 2011; Hahn *et al.*, 2012; Hoffmann *et al.*, 2015; Wei *et al.*, 2019). Genome duplication may in itself be an advantage because it buffers the organism against deleterious alleles (Voigt-Zielinski *et al.*, 2012; Wagner *et al.*, 2019), and higher rates of heterozygosity reduce risks associated with inbreeding effects (Ronfort, 1999). Despite the potential benefits of polyploidy, there are known disadvantages, including the potential dilution of beneficial mutations (Stebbins, 1971) and disturbance of cellular functions such as epigenetic regulation, mitosis, and meiosis (Comai, 2005). However, ploidy polymorphism may provide an important evolutionary pathway for species to establish in previously unsuitable habitats or adapt *in situ* (Grabowski *et al.*, 2014).

Understanding patterns of genetic diversity and evolutionary mechanisms for adapting to new environments is key to improving the conservation of intact grasslands and the restoration of degraded grassland habitats. Globally, restoration practices largely advocate the use of seed sourced from local provenances, based on the assumption that local genotypes are best matched to stable local environments and to avoid perceived risks associated with outbreeding (Thornhill, 1993; Edmands, 2006). Yet, in many cases local provenancing can lead to poor restoration outcomes (Broadhurst *et al.*, 2008; Prober *et al.*, 2015). In highly modified landscapes the genetic integrity of many species has been compromised, and local-provenancing can favour the selection of genetically depauperate and maladapted seed (Jones, 2013). Also, local-provenancing gives little consideration to the persistence of plantings under future climates, with growing evidence that genotypes from non-local sources may outperform those sourced locally (Hoffmann *et al.*, 2015; Prober *et al.*, 2015; Breed *et al.*, 2019). In addition, foundation species are especially important during the restoration process because their genetic variation can shape the networks of ecological interaction influencing community assembly, stability, and evolution (Gibson *et al.*, 2012; Lau *et al.*, 2016). Empirically derived restoration strategies are now being widely adopted around the world to support biodiversity, evolutionary potential, and restoration success, and similar approaches should also be employed for ploidy polymorphism.

In this study, we assess patterns of genetic structure, genotype-ploidy-environment associations, and genomic vulnerability in a foundational grassland species. *Themeda triandra*, commonly known as Kangaroo Grass, has a continent wide distribution, is characterised by ploidy polymorphisms (Hayman 1960) and has limited seed dispersal (Everson et al. 2009). The species provides critical ecosystem services supporting grassland habitats throughout Australia, and is widely used in grassland restorations, but is suffering major declines, shows signs of climate stress, and is in need of improved restoration guidelines. Notably, several studies suggest that re-establishment of *T. triandra* is an important first step for the restoration of Australia’s grasslands (Adair & McDougall 1987; McDonald 2000; Cole & Lunt, 2005), highlighting the importance of research geared toward assessing the resilience of remnant populations, and management approaches that incorporate evolutionary potential. In this context, we assess the likely drivers of genetic structure across a portion of *T. triandra*’s range, predicting both isolation-by-distance (IBD) and isolation-by-environment (IBE) to be key drivers due to the species’ limited seed dispersal and broad climatic niche. Based on estimates of gene flow and correlative measures of local adaptation, we test for genomic mismatches between local gene pools and future climates to help identify populations likely to be most vulnerable to new climatic challenges. Lastly, we test for associations between polyploidy and harsh climate zones, to gain insights into the role of polyploidy in historical and future adaptive processes. These results will provide clear pathways on how to incorporate genomic, environmental, and ploidy information into improved guidance for adaptive management plans that aim to protect these dwindling grassland ecosystems.

## Materials and Methods

### Species and sampling

*Themeda triandra* is a perennial C4 tussock grass, with ploidy variability, and occurs across three continents (Australia, Asia, and Africa) (Dell’Acqua *et al.*, 2013; Snyman *et al.*, 2013; Linder *et al.*, 2018). It is Australia’s most widespread species, being adapted to habitats as diverse as the semi-arid interior and sub-alpine regions (Mitchell & Miller, 1990). In Australia, diploids and tetraploids are the most common ploidy variants, but triploid, pentaploid, hexaploid and aneuploid individuals have also been identified (Hayman, 1960). Past studies suggest that *T. triandra* originally evolved in tropical Asia and migrated through coastal corridors to Australia (Hayman, 1960), with Australian lineages diverging 1.37 mya (0.79 - 3.07 mya) (Dunning *et al.*, 2017). However, dating using secondary calibrations, as in (Dunning *et al.*, 2017) can lead to unreliable and overly young estimates of divergence (Schenk, 2016). *Themeda triandra* is widely considered a foundation species for three reasons: 1) it defines particular ecosystems (Snyman *et al.*, 2013), 2) it controls the distribution and abundance of associated flora and fauna (Morgan, 1998), and 3) it regulates the core ecosystem processes especially through fire (Morgan & Lunt, 1999). The species is also considered to be an indicator of (agro)ecosystem health (Novellie & Kraaij, 2010) and its long-term persistence provides ecosystem stability, ecosystem services, resistance to plant invasions, and facilitates rehabilitation of polluted and degraded habitat (Novellie & Kraaij, 2010; Dell’Acqua *et al.*, 2013). Furthermore, its persistence is critical for the restoration of grasslands in Australia and is reliant on recurring fire to remove old tillers and for seedling establishment (McDougall 1989). The distribution of *T. triandra* is suggestive of a complex evolutionary history with high levels of genetic structuring throughout Australia. Although *T. triandra* itself is not formally listed as an endangered species, it is an important constituent of temperate grassland communities, which have been declared as endangered in the Australian Capital Territory and New South Wales, and threatened in Victoria. The grasslands are under threat due to loss and fragmentation of habitats through inadequate land management practices.

Samples were collected between 2015 and 2017 from 52 populations spanning the heterogeneous climate from its eastern Australian distribution, which deliberately coincides with the densest portion of its distribution. Sampling was structured to ensure different environment combinations were sampled between coastal and inland (west of the Great Dividing Range, see Fig S1) sites. Sites were identified using records on the Atlas of Living Australia public database (ala.org.au) and chosen using the following criteria: herbaria collection or observation was after the year 2000, location data was within 50 m of accuracy, and occurred on land that was publicly accessible. Between 10 and 21 leaf samples were collected per location and placed directly into silica gel to rapidly dessicate leaf samples for DNA preservation. Sampled plants were at least 5 m apart to ensure independence of genotypes by minimising the chance of collecting clonal samples. Our collections comprised a total of 584 individual specimens, which were stored under laboratory conditions until required for genetic analysis.

Using the work of Hayman (1960), we created a predictive map of ploidy levels for populations distributed across our sampling distribution. Hayman measured ploidy levels across Australia, with most of his sites overlapping our sampling distribution. We interpolated his data using nearest neighbor analysis using QGIS v2.14 (Quantum GIS Development team), allowing us to extract predicted ploidy level for each population location to provide us with the number of predicted chromosomes (i.e. diploid = 20; tetraploid = 40; hexaploid = 60). A few individuals were equidistant between two predicted ploidy levels and were assigned ploidy level between 20 and 40. This was interpreted as indicating a mixed ploidy population. Ploidy predictions were verified with population-level heterozygosity, see below for details.

### DNA extraction and library preparation

For reduced-representation library preparation and sequencing, genomic DNA from each individual was isolated from approximately 25 mg of silica-dried leaf tissue using the Stratec Invisorb DNA Plant HTS 96 kit (Invitek, Berlin, Germany). Libraries were created similarly to Ahrens et al. (2017). Briefly, extracted DNA was digested with PstI for genome complexity reduction, and ligated with a uniquely barcoded sequencing adapter pair. We then amplified each sample individually by PCR to avoid sample bias. We pooled samples in equimolar ratios and selected amplicons between 350 and 600 bp from an agarose gel. The library pool was sequenced on three Illumina NextSeq400 lanes using a 75bp paired-end protocol on a high output flowcell at the Biomolecular Resources Facility at the Australian National University, generating ~864 million read pairs.

For long-reads via the MinION sequencer (Oxford Nanopore Technologies, UK), we used the open access high molecular weight DNA extraction protocol developed by Jones & Borevitz (2019). Briefly, 30 g of fresh leaf material from a known diploid individual was processed with 150 mL nuclei isolation buffer using a high-powered blender. The homogenate was filtered repeatedly using a funnel, through sequentially 2, 4 and 8 layers of Miracloth. Next, 100% Triton X-100 was added for nuclei isolation and the mixture centrifuged to create a pellet of nuclei. The pellet was washed twice with a pre-chilled nuclei buffer. DNA extraction from the nuclei was initiated by adding fresh lysis buffer with 3% Sodium dodecyl sulfate (SDS) at 50ºC. Binding buffer was added to use Sera-Mag beads to remove the lysis buffer from the DNA solution, washing with 70% ethanol 3 times until the beads were clean. The beads were removed by adding 220 uL of ultra-pure H_2_0 and resuspending the beads with attached DNA. The supernatant was removed and subsequently size selected for fragments longer than 30 kb using a PippinHT (Sage Science, Beverly MA). MinION library preparation and sequencing was performed as per the manufacturer’s instructions and specifications, and resulted in 412,906 reads (Fig S2). Median read length was 27,156 bases, and the longest read length was 144,466 bases, with an overall average read-quality of 10 (Fig S2).

### SNP calling

We checked the quality of the raw short-read sequencing reads with FastQC (v0.10.1, [Andrews, 2012]). Then, we demultiplexed the raw reads associated with each sample’s unique combinatorial barcode using AXE v0.2.6 (Murray & Borevitz, 2018). During this step we were unable to assign 19% of the reads. We trimmed each sequence to 64 basepairs while removing the barcodes and ensured quality of the reads using trimmomatic v 0.38 (Bolger *et al.*, 2014). Quality was assessed using a sliding window of 4 basepairs (the number of bases used to average quality) and a quality score of 15 (the average quality required among the sliding window), and if the average quality dropped below 15, the sequences were cut. Then we indexed the long-reads (Fig S2 for distribution of length and number of reads sequenced) using the BWA software and the *index* argument. We aligned the short-reads to the long-reads for more accurate SNP calling compared to a *de novo* pipeline. Short-reads were aligned using BWA-mem (v0.7.17-r1198, [Li *et al.*, 2013]), as paired reads, with 82.5% of reads successfully mapped. Samtools v 1.9 (Li *et al.*, 2009) was used to transform the SAM files to BAM files for use within STACKS v 2.41 (Catchen *et al.*, 2013). The argument gstacks and populations were used in that order on the BAM files to create a VCF file, minimum thresholds (minor allele frequency = 0.01; one random SNP per read was retained) were set here for further cleaning in R (R core development team 2019). The mean coverage per sample was 15.8× with a standard deviation of 20×, this resulted in many samples being dropped (see below for details). Lastly, VCFtools v 0.1.16 (Danecek *et al.*, 2011) was used to create a 012 file for further cleaning of the snp matrix in R.

The missing data threshold was set to 50% per locus and individual which resulted in an average of 30% missing data from the whole SNP dataframe. Minor allele frequency was set to 0.05 to avoid identifying patterns of population structure that may be due to locally shared alleles (De la Cruz & Raska, 2014). Then we removed SNPs in high linkage disequilibrium (>50% similar). We also removed possible clones in Genodive v 2.0b27 (Meirmans & Van Tienderen, 2004) using the *assign clones* function, removing nine individuals. After conservative SNP filtering, we were left with 487 individuals from 52 populations.

### Analysis

Genodive was used to estimate population summary statistics for the total number of alleles observed across loci, total heterozygosity, and the inbreeding coefficient (*G*_IS_; Nei, 1987). We expected that the degree of heterozygosity within populations would reflect ploidy status (i.e. higher heterozygosity would imply polyploids) as described by Soltis & Soltis (2000). Consequently, we validated predicted ploidy level among populations from Hayman’s map (see above for details) by comparing those predictions to population-level heterozygosity. *G*_IS_ is the same as *F*_IS_ for a single locus with two alleles (Chakraborty & Leimar 1987), and is calculated by the ratio of observed heterozygosity within subpopulations to the expected heterozygosity and ranges from −1 (complete outbreeding) to 1 (complete inbreeding). Genodive was also used for an analysis of molecular variance (AMOVA) using the Excoffier method (Excoffier et al. 1995). Global *F*_ST_ with 95% confidence intervals was calculated using the *fstat* argument and the population pairwise *F*_ST_ was calculated using the *pairwise.fst* argument in the *hierfstat* package in R (Goudet, 2005).

*Themeda triandra* has a broad geographic distribution spanning a variety of environmental gradients, therefore we wanted to estimate the amount of genetic variation that could be attributed to isolation-by-distance (IBD) and -environment (IBE). First, we downloaded the 19 bioclim variables from worldclim.org (Fick & Hijmans, 2017), and extracted all of the climate variables for each of the sample locations in R using the package *rast*er (Hijmans & van Etten 2012). A Principle Components Analysis (PCA) was performed to determine potential correlations between the 19 climate variables and produce an environmental dataset consisting of least correlated variables (Fig S3). We chose to retain variables from six of the loose clusters (temperature mean diurnal range (T_RANGE_), maximum temperature of the warmest month (T_MAX_), precipitation seasonality (P_SEAS_), mean annual temperature (T_MA_), mean annual precipitation (P_MA_), and precipitation of the driest month (P_DM_)).

We used sNMF (Frichot *et al.*, 2014) in the LEA package in R (Frichot & François, 2015) to investigate the observed patterns of population structure that include contributions from both geography (IBD) and environment (IBE). sNMF estimates ancestry coefficients based on sparse non-negative matrix factorisation and least-squares optimisation. The sparse non-negative matrix factorisation is robust to departures from traditional population genetic model assumptions, making this algorithm ideal to use with polyploid species such as *T. triandra*. We performed sNMF with the following attributes: *k* = 1-10, 10 replications per *k*-value (number of ancestral clusters), and 1,000 iterations. Entropy scores for each *k*-value were compared to choose the optimal number of clusters using the recommendations in the sNMF instruction manual. A consensus for the optimal *k*-value was created by averaging the results over the 10 replicate runs using CLUMPP v1.1.2 (Jakobsson & Rosenberg, 2007) and drawn using DISTRUCT v1.1 (Rosenberg, 2003).

We used Moran’s Eigenvector Maps (MEM) to test if IBD was a major determinant of the species’ genetic diversity, as described in previous work (Dray *et al.*, 2006; Legendre & Legendre, 2012) but called PCNM in the first papers. Briefly, MEM calculates a matrix of pairwise Euclidean distances **D** among the sampling sites, then transform the **D** matrix into a similarity matrix to produce the MEM. Eigenvalues are produced corresponding to orthogonal vectors of similarity. To ascertain spatial patterns of genetic diversity we used the R package memgene (Galpern *et al.*, 2014). Memgene identifies spatial neighbourhoods in genetic distance data that adopts a regression framework where the predictors are generated using MEMs, this multivariate technique was developed for spatial ecological analyses but is recommended for genetic applications. Memgene identifies variables (eigenvalues) that represent significant spatial genetic patterns at multiple spatial scales. Each variable explains a proportion of the total variance explained by spatial patterns. For this study, we show two variables because it explains most of the variation described by IBD.

Using the environmental data layers we employ a generalized dissimilarity model (GDM) to identify the importance of specific climate variables responsible for shaping observed patterns of genetic structure within our dataset. Analyses were performed using the gdm package v 1.3.7 in R (Manion *et al.*, 2018) and a pairwise *F*_ST_ matrix (based on all SNP loci) to estimate allelic turnover through climatic space (deviations in allele frequency associated with environment type). Where GDM holds all variables in the model constant to identify the partial genomic distance associated with the climate factor (Ferrier *et al.*, 2007), whereby accounting for spatial patterns caused by demographic processes (Fitzpatrick & Keller, 2015). After running the GDM analysis, only four of the climates remained (T_MAX_, P_SEAS_, T_MA_, and P_MA_), as the other two climate factors were removed by a backward□elimination procedure. The GDM output includes the deviance explained by the climate and spatial variables, and a spline plot for each climate and spatial variable. Spline plots were predicted across the study area and beyond for every 2.5km grid cell. These predicted grids were mapped using ggplot in R (Wickham, 2011) to describe the relative IBE.

We calculated ‘genomic vulnerability’ for the sampling area following Bay et al. (2018), which consists of three main components: exposure, sensitivity, and adaptive capacity (Dawson *et al.*, 2011). Genomic vulnerability is the amount of genomic change required to track environmental change over time and is interpreted as expected population decline. To do this, we substituted predictive maps in 2070 using the CCSM4 model with the representative concentration pathway 8.5 (worldclim.org), which is a prediction based on the anthropogenic carbon dioxide output not deviating from its current trajectory. These maps were also downloaded from worldclim and developed in the same way as described above. Lastly, we subtracted the projected genomic differentiation from the current genomic differentiation to get a difference between the two. We estimate genomic vulnerability twice, with and without predicted ploidy levels to understand how ploidy may affect population decline, particularly in the most vulnerable areas.

## Results

We estimated patterns of population structure among 487 samples from 52 sample locations for *T. triandra* using a dataset consisting of 3,443 polymorphic SNPs with a minor allele frequency (MAF) of 0.05 and an average of 30% missing data. AMOVA indicated that a significant proportion of the genetic variance (10%) could be attributed to difference among sample sites (*P* = 0.001; *F*_ST_= 0.22), while the majority of the variance (79.3%) was attributed to differences between individuals (*P* < 0.01; *F*_IT_= 0.31). Large and significant positive inbreeding coefficients (*G*_IS_) were observed for many sites, indicating an excess of homozygotes, while three populations had negative inbreeding coefficients indicating homozygote deficits (Table 1). Levels of genetic diversity (number of alleles and heterozygosity) was variable among populations, with a mean number of alleles of 1.082 (95% CI 1.078-1.086; range 1.109 - 1.366) and a mean heterozygosity within populations (H_S_) of 0.074 (range 0.06 - 0.12; Table 1). Heterozygosity estimates reflect patterns that are consistent with the hypothesis that greater ploidy levels are present in the hotter regions of our sampling distribution (Fig 1). However, this linear model, although significant (r^2^= 0.086; *P* = 0.035), explains only a small proportion of the variation. This pattern is likely driven by the three populations in the hottest region. Heterozygosity and predicted chromosome number were in agreeance for these three populations, the populations with the highest T_MAX_ (QLD, PR, SWC). Some populations with high heterozygosity were predicted to be diploids (UL, GOR, NAM), but these populations were nearly equidistant to tetraploid and diploid populations and are likely tetraploid populations (Fig 1).

**Table 1.** Locations and genetic diversity indices for sampled populations. T_MAX_ = maximum temperature of the warmest month; P_SEAS_ = precipitation seasonality; H_s_ = heterozygosity within populations; G_IS_ = inbreeding coefficient; A_N_ = number of alleles; C_P_ = predicted chromosome number.

| Pop | X | Y | $T_{MAX}$ (°C) | $P_{SEAS}$ (mm) | $A_N$ | $H_s$ | $C_P$ | $G_{IS}$ |
| --- | --- | --- | --- | --- | --- | --- | --- | --- |
| MTG | 138.754 | -34.977 | 26.7 | 52 | 1.357 | 0.088 | 40 | 0.059 |
| BBNP | 153.028 | -30.420 | 28.1 | 43 | 1.215 | 0.066 | 20 | 0.181 |
| BCR | 153.054 | -28.646 | 28.2 | 45 | 1.241 | 0.071 | 20 | 0.182 |
| BL | 151.737 | -29.867 | 25.6 | 33 | 1.233 | 0.068 | 20 | 0.2 |
| BLAPT | 152.807 | -31.395 | 26.9 | 34 | 1.232 | 0.079 | 20 | 0.317 |
| BLARD | 150.444 | -35.196 | 25.3 | 24 | 1.21 | 0.066 | 20 | 0.154 |
| BNR | 151.997 | -29.113 | 25.2 | 32 | 1.169 | 0.064 | 20 | 0.156 |
| BRA | 151.996 | -32.631 | 27.2 | 25 | 1.212 | 0.064 | 20 | 0.179 |
| BU | 151.076 | -30.189 | 29.5 | 31 | 1.164 | 0.064 | 20 | 0.207 |
| BYR | 153.620 | -28.652 | 28.1 | 32 | 1.203 | 0.064 | 20 | 0.143 |
| CB | 150.674 | -30.885 | 30.9 | 34 | 1.164 | 0.062 | 30 | 0.137 |
| BCG | 143.316 | -37.612 | 26.1 | 23 | 1.142 | 0.043 | 20 | 0.085 |
| DCD | 150.728 | -34.013 | 28.1 | 29 | 1.294 | 0.083 | 40 | 0.121 |
| DCR | 149.982 | -36.356 | 24.8 | 24 | 1.271 | 0.08 | 30 | 0.062 |
| DW | 151.997 | -29.114 | 25.2 | 32 | 1.203 | 0.068 | 20 | 0.165 |
| RWCK | 146.817 | -36.583 | 28.1 | 32 | 1.185 | 0.062 | 22 | 0.366 |
| BUR | 145.026 | -37.834 | 26.0 | 17 | 1.136 | 0.06 | 20 | 0.328 |
| ANG | 144.153 | -38.335 | 23.9 | 22 | 1.187 | 0.07 | 20 | 0.138 |
| EUN | 152.888 | -30.811 | 27.7 | 39 | 1.234 | 0.067 | 20 | 0.184 |
| GHK | 149.863 | -36.979 | 24.0 | 18 | 1.259 | 0.073 | 20 | 0.236 |
| GOR | 150.588 | -35.009 | 25.2 | 23 | 1.397 | 0.102 | 20 | -0.04 |
| GRES | 151.219 | -32.546 | 30.1 | 34 | 1.231 | 0.068 | 20 | 0.171 |
| QLD | 149.878 | -27.926 | 33.5 | 29 | 1.309 | 0.12 | 40 | 0.034 |
| JG | 152.008 | -30.514 | 25.3 | 38 | 1.187 | 0.065 | 20 | 0.196 |
| KCK | 152.579 | -31.795 | 27.5 | 37 | 1.238 | 0.068 | 20 | 0.177 |
| KOZ | 148.402 | -35.889 | 22.4 | 29 | 1.255 | 0.075 | 20 | 0.28 |
| KUN | 152.844 | -31.196 | 27.3 | 37 | 1.20 | 0.069 | 20 | 0.205 |
| L | 152.292 | -28.411 | 27.9 | 37 | 1.195 | 0.073 | 20 | 0.17 |
| LO | 149.998 | -33.167 | 26.4 | 20 | 1.203 | 0.082 | 20 | 0.318 |
| MGR | 149.077 | -36.244 | 25.0 | 19 | 1.254 | 0.07 | 20 | 0.051 |
| ML | 152.473 | -28.380 | 26.9 | 40 | 1.285 | 0.083 | 20 | 0.218 |
| MNP | 150.373 | -35.457 | 24.1 | 13 | 1.36 | 0.089 | 20 | 0.127 |
| Mong | 149.944 | -35.426 | 25.5 | 16 | 1.193 | 0.064 | 20 | 0.252 |
| MS | 150.881 | -29.988 | 29.8 | 31 | 1.161 | 0.067 | 40 | 0.165 |
| MSF | 149.055 | -34.825 | 28.1 | 13 | 1.293 | 0.084 | 20 | 0.334 |
| NAB | 152.370 | -32.086 | 27.7 | 35 | 1.217 | 0.063 | 20 | 0.226 |
| NAM | 152.976 | -30.639 | 28.0 | 41 | 1.109 | 0.109 | 20 | --- |
| OPC | 153.037 | -29.820 | 28.5 | 40 | 1.109 | 0.064 | 20 | 0.116 |
| PR | 150.186 | -31.418 | 31.7 | 34 | 1.366 | 0.115 | 40 | -0.135 |
| MSCP | 140.631 | -37.145 | 28.1 | 44 | 1.242 | 0.072 | 20 | 0.257 |
| SIW | 153.146 | -30.192 | 27.4 | 40 | 1.147 | 0.062 | 20 | 0.17 |
| SOM | 151.286 | -33.404 | 26.1 | 31 | 1.248 | 0.065 | 20 | 0.187 |
| SPNR | 149.747 | -37.557 | 22.6 | 12 | 1.247 | 0.07 | 20 | 0.157 |
| STCK | 149.314 | -35.360 | 26.7 | 13 | 1.257 | 0.075 | 20 | 0.245 |
| SWC | 149.707 | -31.400 | 31.3 | 31 | 1.18 | 0.105 | 41 | -0.753 |
| NSW | 148.142 | -36.542 | 26.6 | 17 | 1.245 | 0.067 | 20 | 0.151 |
| TOO | 152.391 | -28.453 | 28.7 | 39 | 1.221 | 0.068 | 20 | 0.183 |
| UL | 152.073 | -30.537 | 25.2 | 39 | 1.234 | 0.099 | 20 | -0.326 |
| WOL | 150.806 | -34.434 | 25.6 | 31 | 1.229 | 0.065 | 20 | 0.183 |
| WYE | 151.491 | -33.175 | 26.6 | 30 | 1.235 | 0.065 | 20 | 0.195 |
| YNGNP | 150.693 | -33.057 | 28.2 | 38 | 1.223 | 0.063 | 26 | 0.196 |
| YNR | 151.076 | -30.189 | 29.5 | 31 | 1.213 | 0.076 | 20 | 0.319 |
| Overall |  |  |  |  | 1.842 | 0.074 |  | 0.134 |

**Figure 1.**
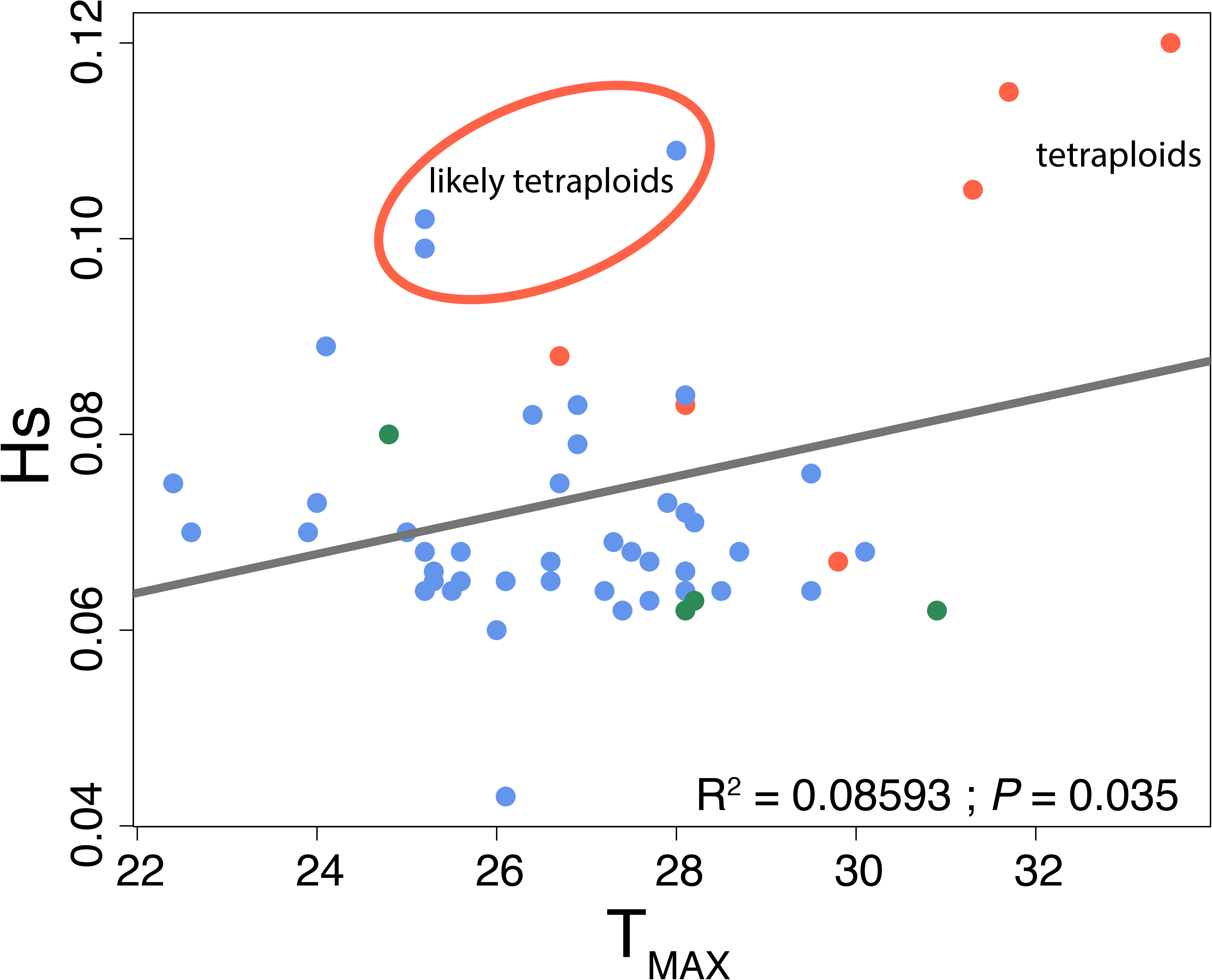
Within population heterozygosity (Hs) versus maximum temperature of the warmest month. Colors indicate diploid (blue), mixed populations (green; equidistant between tetraploid and diploid populations), and tetraploid (red) based on Hayman’s (1960) work. Ellipsoid outlines populations that have high heterozygosity and may be tetraploids.

General patterns of population structure show a clear delineation between southern and northern populations (Fig 2) with an optimal *k*-value of 3 (Fig S4). The third *k*-value is found in two populations, and partially assigned in two other populations. These populations containing the third ancestral cluster were generally found in the central area of the sampling region. Notably, there are portions of populations, particularly in the south central portion of the sampling region, that have been assigned to the northern ancestral cluster. While there are a few individuals in the north assigned to the southern ancestral cluster.

**Figure 2.**
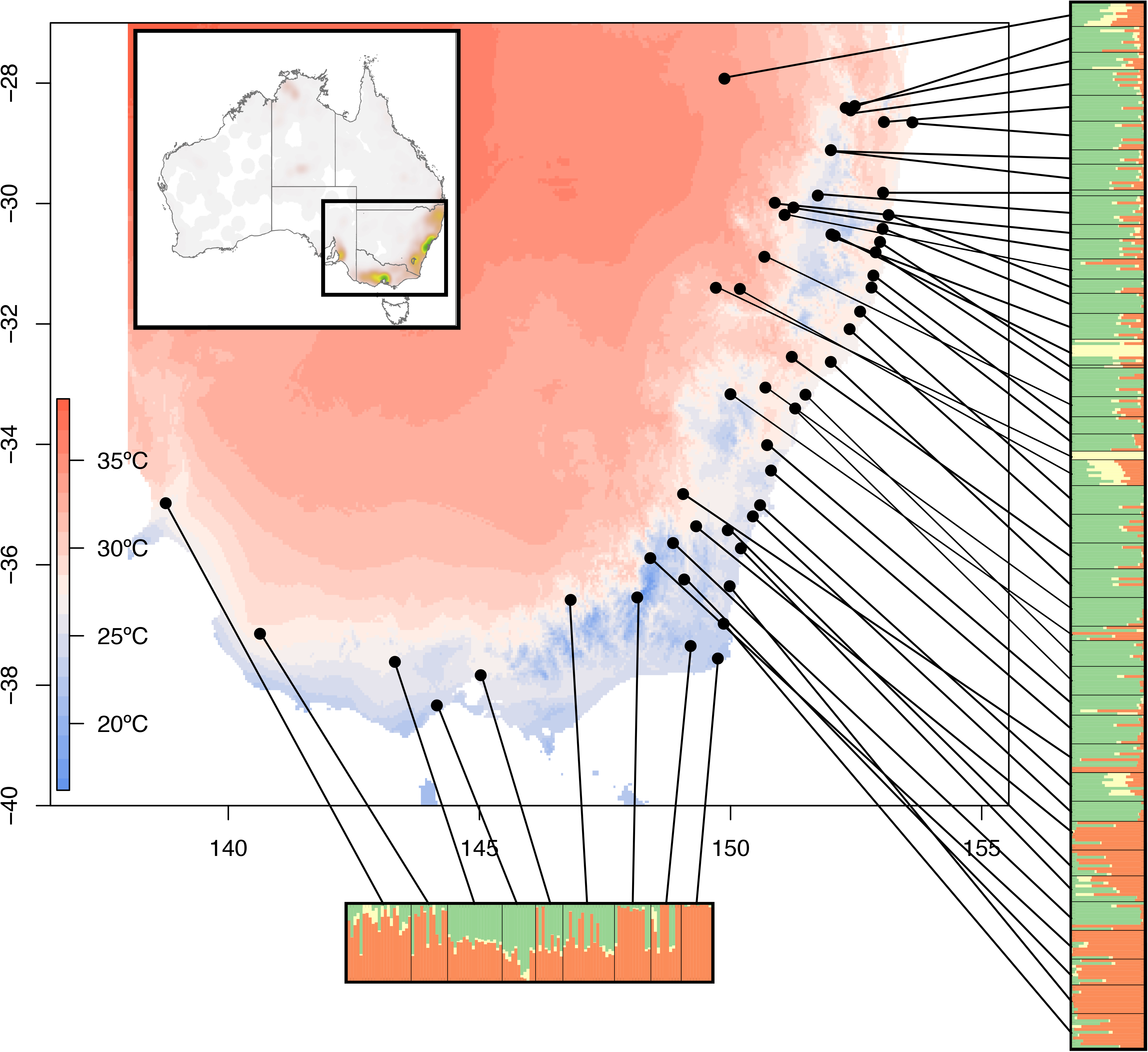
Sparse non-negative matrix factorization (sNMF) for all individuals, points on the map indicate population location, map colors represent T_MAX_ (maximum temperature of the warmest month). Barplot indicates identified genetic ancestral clusters for each individual (bar) with an optimal *k*-value of three. Inset shows the Australia-wide distribution of *T. triandra* as a heat map and location of the study area.

Isolation-by-distance (IBD) was found to be significant in *T. triandra*. In fact, IBD accounts for 54% of the total genomic variation (Fig 3). Two axes are shown in separate figures, and together they explained 95% of the variation explained by IBD alone. The first axis shows a strong split between the northern and southern sections of the sampling area (Fig 3a), similar to the population structure identified in the sNMF results. A second pattern of IBD occurs in the northern part of the sampling region and is between the inland and coastal populations, while the most westerly population is slightly more similar to the northern sampling region (Fig 3b).

**Figure 3.**
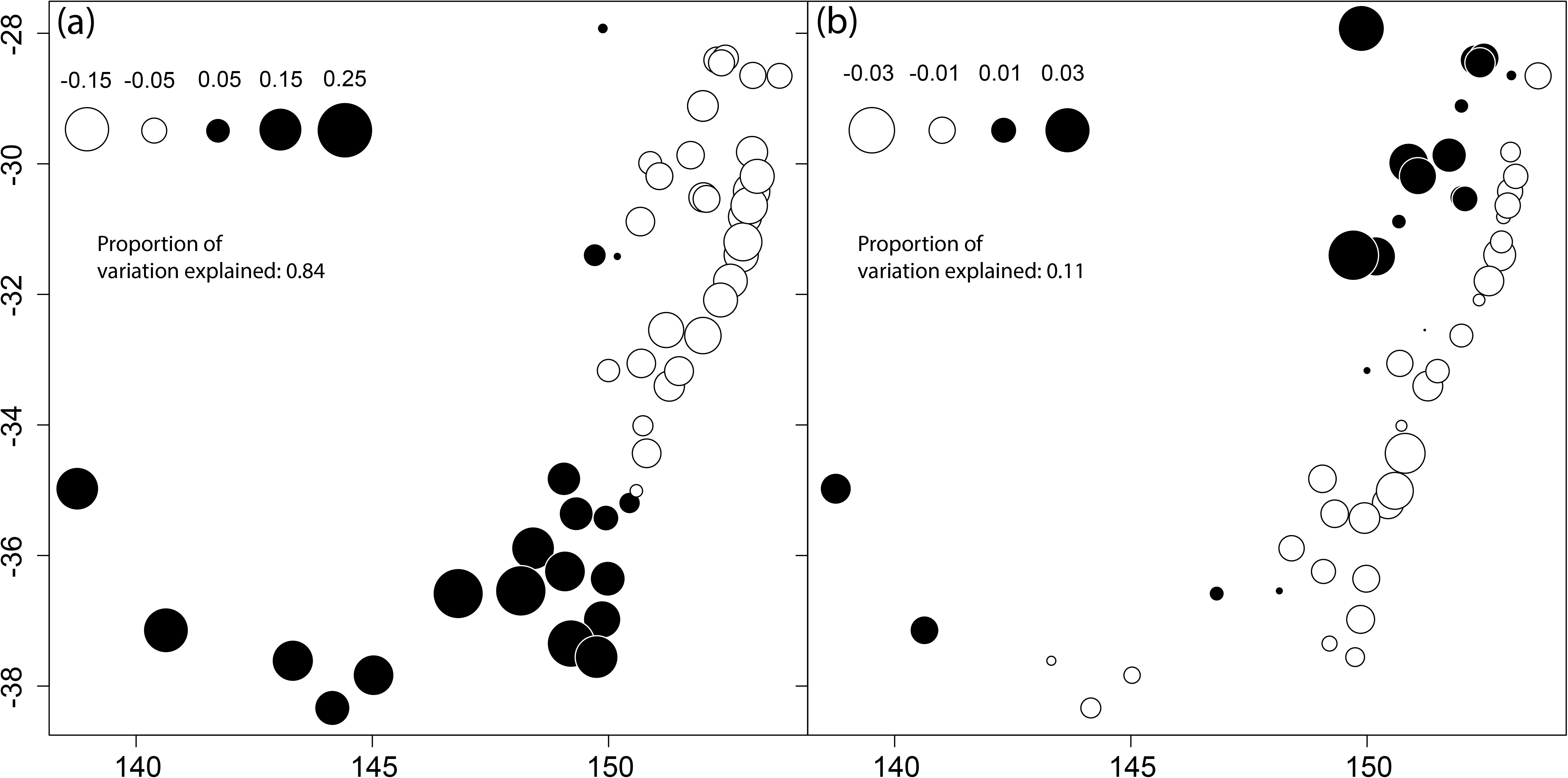
Identification of the spatial component of genetic variation using Moran’s Eigenvector Maps. Two distinct spatial patterns accounted for most of the 54% of genetic variation explained through isolation by distance. The first MEM variable (a) explained a greater proportion of the variation than the second variable (b). Circles of similar size and colour represent individuals with similar scores on this axis.

In addition to spatially driven genomic variation, isolation-by-environment (IBE) explains a significant amount of variation. While we chose six independent climate variables to explore IBE, only four were found to be significant (T_MA_, T_MAX_, P_MA_, P_SEAS_; maps for climate variables in Fig S5). The GDM analysis was able to identify that 31.3% of the variation was attributable to these climate and spatial variables (Fig 4), and 22.0% of the variation was attributable directly to climate. When performing the same analysis with the inclusion of ploidy level, the variation explained rose by only 0.4%, but under this model, the T_MAX_ variable explained less variation (red lines in Fig 4) while all other variables remained similar. In the current climate, the differences between the two models were negligible (Fig 5a & c). However, when forecasting the differences in 2070, the outputs suggest a heterogeneous population decline by 0 and 25% (Fig 5b) with the highest proportion of change occurring inland of the eastern coast. Critically, the inclusion of ploidy polymorphism showed genomic vulnerability dropping by 5% in the most vulnerable areas (Fig 5b & d; ploidy map provided in Fig S5), in this output, we find that genomic vulnerability occurs where the land transitions from the alpine region to the inland region. The lowest probability of change (population decline or gene pool turnover) is in the mountainous ecosystems in the southeastern portion of the sampling region.

**Figure 4.**
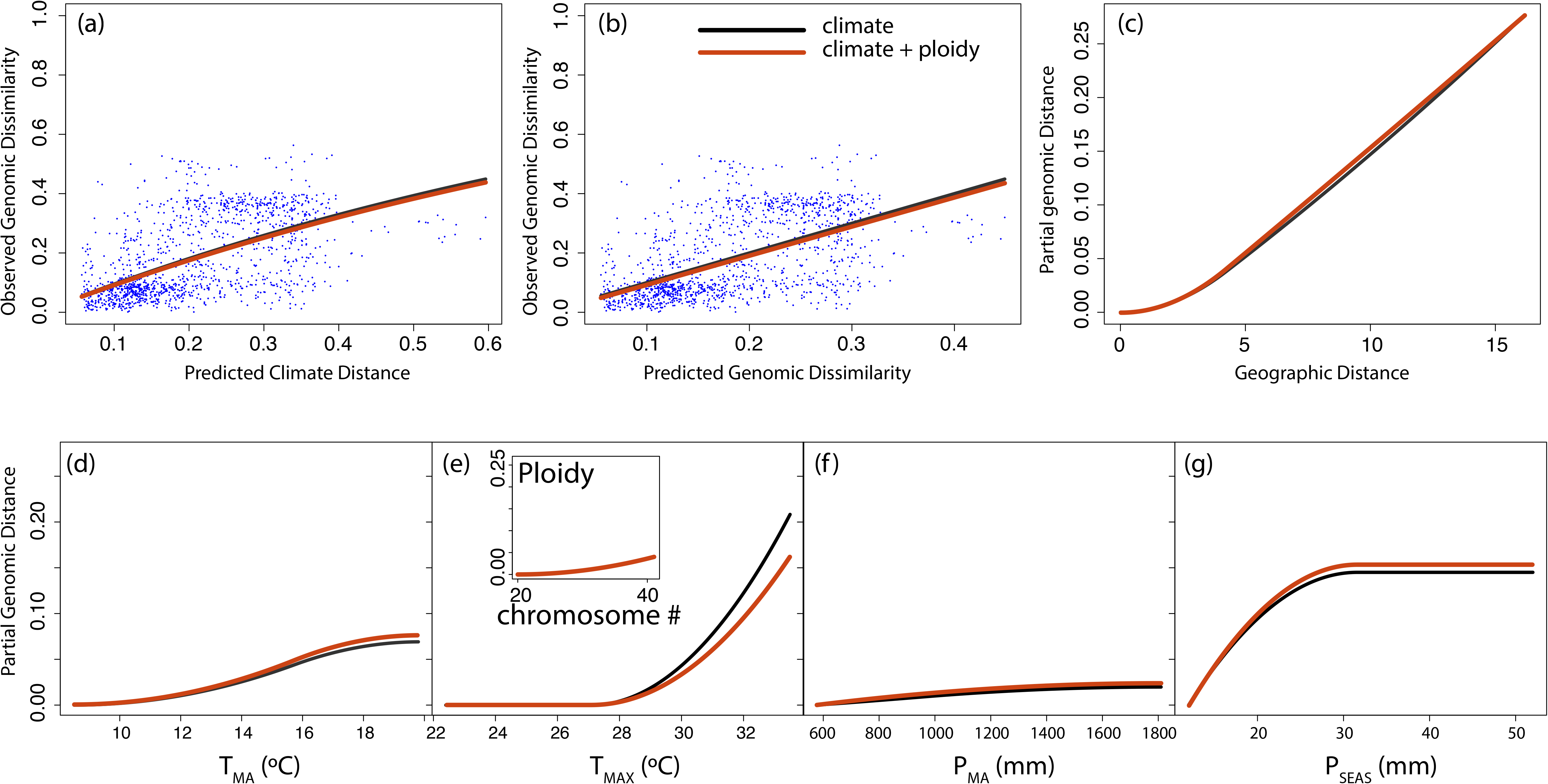
Generalised dissimilarity modelling (GDM). (a) Non-linear relationship between climate distance and genomic distance, where points are site pairs. (b) Relationship between predicted genomic distance and observed genomic distance, where points are site pairs. (c) The geographic spline showing the relationship between predicted genomic change and geographic distance. (d–g) Predicted splines showing the estimated relationship between genomic distance and individual climate variables: (d) mean annual precipitation (T_MA_), (e) maximum temperature of the warmest month (T_MAX_), (f) mean annual precipitation (P_MA_), and (g) precipitation seasonality (P_SEAS_); inset is the amount of variation explained by predicted ploidy polymorphisms (red lines are the model that includes ploidy). Variation explained for the climate-only + spatial model is 31.3% (22% attributed to climate), and with climate, ploidy, and spatial is 31.7% (23% attributed to climate).

**Figure 5.**
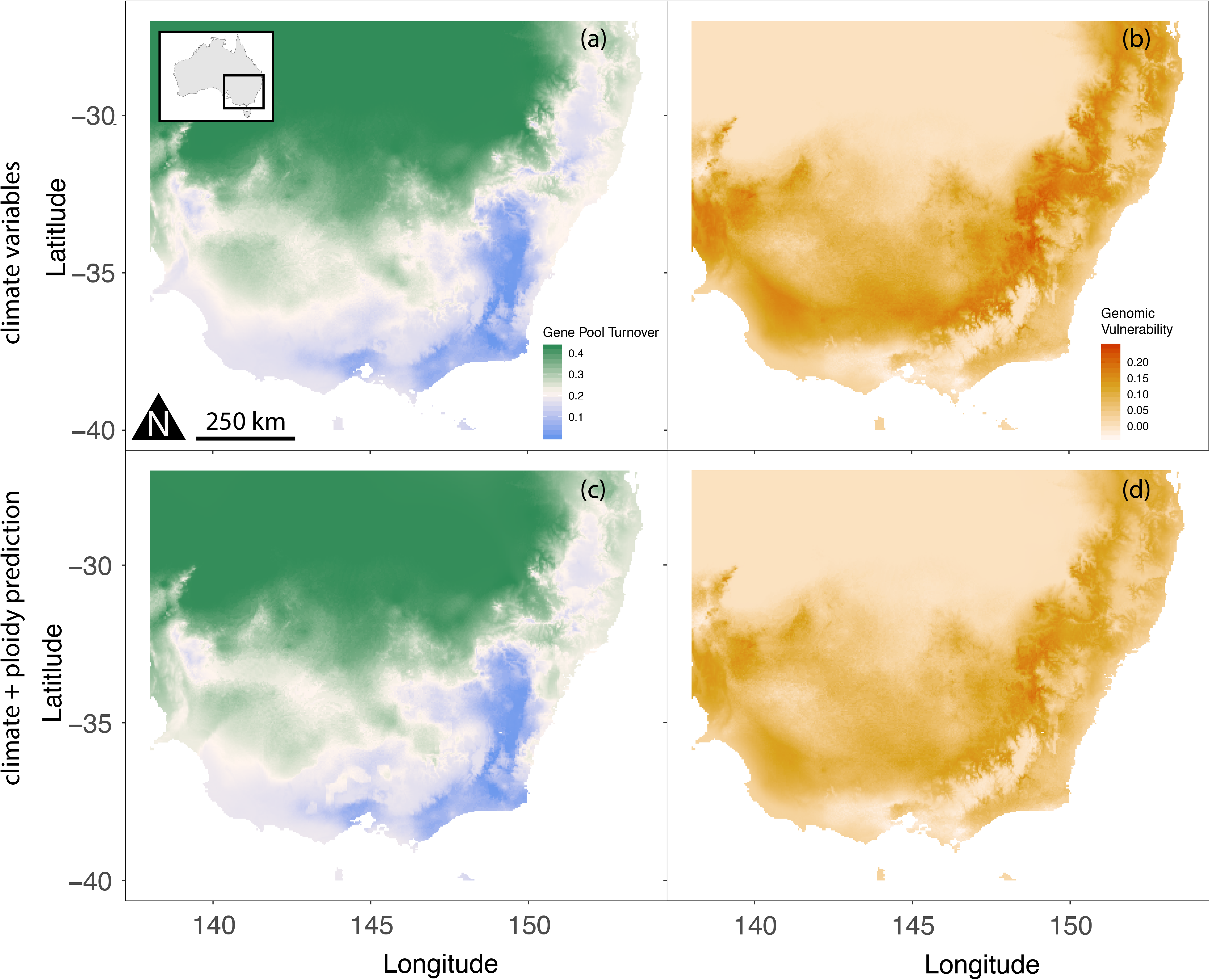
Predicted spatial variation in genomic composition based on the outputs from the general dissimilarity models (GDM). Maps include the (a) climate-only GDM and (b) the predicted genomic vulnerability based on comparing the current GDM and the predicted GDM for 2070. Whereas, the (c) climate + ploidy GDM, and (d) and the predicted genomic vulnerability are shown for direct comparison to the climate-only model. A 5% reduction in genomic vulnerability is indicated in the most severely affected areas when including ploidy level in the GDM. The greater the difference (dark orange), the more genomic change is needed to adjust to future climate conditions.

## Discussion

Our study indicates contemporary structuring of genomic diversity in *Themeda triandra* is being driven largely by a combination of spatial and climate factors. These patterns are indicative of a species with limited propagule dispersal and restricted gene flow. The apparent lack of connectivity among remnant populations suggests gene flow is unlikely to help local populations adapt to future climate challenges. Instead, their adaptive potential will rely on trait plasticity and standing genetic variation that allows for adaptation *in situ*. Strong associations between gene pools and climate may reflect patterns of local adaptation, and heterogeneity in climatic conditions at both local and regional scales, suggests that the impacts of climate change on remnant populations are likely to be uneven. This is supported by assessments of mismatches between current and predicted future genomic variation, creating heterogeneous patterns of ‘genomic vulnerability’ across populations. We also demonstrate polyploidy associations with harsh climate zones, suggesting polyploidy is potentially linked to historical adaptation processes and may assist populations in overcoming future climate challenges. This study highlights the need for adaptive management strategies that incorporate evolutionary potential, including seed sourcing and population mixing strategies that can help overcome genomic vulnerability and maladaptation under future climates.

### Isolation-by-distance

The majority of genomic variation found in *T. triandra* could be explained by geographic isolation. This is likely to be due to low levels of gene flow and seed dispersal between populations contributing to strong genetic structuring, as found in South African populations (Everson *et al.*, 2009). However, this structure could also be driven by a partially apomictic reproductive system in *T. triandra* (Brown & Emery, 1957; Birari, 1980), with clonal reproduction inflating signals of population-level genetic uniqueness. We found some evidence of clonal *T. triandra* genotypes, but these individuals were removed during the data filtering phase prior to analyses. While our data are unable to confirm the relationship between clonality and polyploidy due to low replication, our data suggests that polyploidy occurs infrequently at milder temperatures, while being dominant among populations occurring in the highest temperature environments. These findings are consistent with Hayman (1960) who argues that the diploid landrace is likely absent in the harsher climates, suggesting the presence of positive selection for polyploid landraces in the hot and dry inland environments.

Perhaps the most germane work of this nature is that of the grass species *Panicum virgatum*. Similar to *T. triandra*, *P. virgatum*’s ploidy level increases with distance from the coast, with higher ploidy levels found in more arid inland environments (Zhang *et al.*, 2011; Lowry *et al.*, 2014; Grabowski *et al.*, 2014). As demonstrated in *P. virgatum,* we provide evidence for polyploidy evolution through multiple, isolated events rather than the establishment and expansion of polyploids from one duplication event. For example, some populations of predicted polyploids are more closely related to diploid populations rather than other tetraploid populations. This suggests genome doubling can occur spontaneously within populations and is both induced and maintained by selection under certain environmental scenarios. Indeed, it has been shown that polyploids can have an increased fitness advantage under heat- and water- stressed conditions (Rey *et al.*, 2017).

### Isolation-by-environment and genomic vulnerability

Along with geography, climate factors describe a large percentage of genomic variation found in *T. triandra*. We found strong associations between gene pools and environments (particularly with T_MAX_ and P_SEAS_), possibly reflecting adaptation to climate. While quantitative tests are needed to validate these findings (e.g. common garden experiments – Sork, 2017), our results are consistent with the idea that signals of adaptation are ubiquitous throughout genomes (Kern & Hahn, 2018). Maximum temperature of the warmest month or week (T_MAX_) has been found to be an important driving force of selection in other Australian plants (Steane *et al.*, 2017a,b; Jordan *et al.*, 2017; Ahrens *et al.*, 2019). Interestingly, evidence suggests that climatic factors can have different impacts on patterns of genetic diversity and adaptation in different grass species. For example, *T. triandra* and *Andropogon gerardii* are both dominant C4 grass species, with temperature and precipitation factors being key selective forces driving diversity in *T. triandra,* while lower precipitation suppresses genetic diversity in *A. gerardii* (Avolio *et al.*, 2013). Despite these differences, polyploidy appears dominant in harsher regions in both species indicating there are ploidy based adaptive responses to climate, enabling the expansion of species into habitats unsuitable or less suitable for diploids. The line of adaptation demarcation is stronger for *T. triandra*, where persistence in the semi-arid landscape appears entirely dependent on polyploids, compared to *A. gerardii*, where ploidy mixing occurs in harsher parts of its climate range (Keeler, 1990).

Our analyses of genomic vulnerability across the study area suggest that some populations of *T. triandra* will be more adversely impacted by climate change than others. For example, the most inland populations of our sampling are most vulnerable where we estimate that populations will need to change by over 20%, this region includes both diploid and polyploid populations. The least vulnerable populations are located in the southern and mountainous regions where we would expect populations to change by 0 to 5%. The future mismatch of predicted gene pools in some regions suggests that a change of as much as 25% will be necessary for adaption to the new challenges. Our predictions are based only on correlative analyses, and caution should be taken when interpreting these findings given the uncertainty associated with the genetic mechanisms (i.e. epistatic interactions (Juenger *et al.*, 2005), pleiotropy (Solovieff *et al.*, 2013), chromosomal rearrangements (Juenger *et al.*, 2005; Yeaman, 2013), and polyploidy (Van de Peer *et al.*, 2017)) and ecological interactions likely to dictate future adaptive responses (Fordyce, 2006). Indeed, our findings further highlight the need for quantitative experiments (i.e. common garden) to validate these findings by testing the physiological limits and safety margins of individual populations.

Not surprisingly, the genomic vulnerability of several populations was buffered by as much as 5% by the presence of polyploids, and this is likely to be an underestimation due to under-predicting which populations are polyploids. Polyploidy is known to provide fitness advantages in many plant species persisting in hot and arid environments, including *T. triandra* populations (Godfree et al. 2017). The increased heterozygosity associated with polyploidy may have the effect of slowing the loss of genetic variation and providing more variants for selection to act upon (Comai, 2005). Elevated fitness may also be influenced by duplicated genes and genomes, each set capable of independent selection and evolving new functions (Soltis & Soltis, 2000) by retaining multiple gene copies and acquiring a new function in one copy (Wendel, 2000). Further, increased performance could be due to differential levels of expression between ploidy landraces (e.g. Cromie *et al.*, 2017; Wang *et al.*, 2018; Liqin *et al.*, 2019), and be partially dependent on different epigenetic patterns (Nagymihály *et al.*, 2017). However, quantitative measures are needed to determine how differential expression between diploid and tetraploid landraces may affect their ability to persist in their optimal climates. We argue that these types of processes are likely occurring in *T. triandra* landraces, allowing polyploids to persist and outperform their diploid counterparts in hotter and drier climates.

### Management and restoration implications

We are at a critical juncture in history where management and restoration of grassland ecosystems is necessary to preserve these ecosystems and their services. However, the interplay of habitat fragmentation and rapid climate change poses a significant challenge for the conservation and restoration of functionally important plant species. Prioritising investments requires an understanding of species biology and ecology to apply frameworks for identifying the species and populations most at risk. *Themeda triandra*, the most widely distributed species in Australia, is at a critical inflection point due to its use as a food crop (Pascoe, 2018), for native pasture (Fourie *et al.*, 1985), as a foundational species (Snyman *et al.*, 2013), for selective breeding (e.g. *Lolium*/*Festuca* – Yamada *et al.*, 2005), and in the restoration of degraded lands (Cole & Lunt, 2005; Snyman *et al.*, 2013). Our results provide a critical first step and baseline information to support these new interests, future studies and the development of empirically based management strategies that target grassland and open woodland ecosystems. In Australia, research efforts have mostly focused on *Eucalyptus* species, finding that eucalypt populations are often connected by high levels of gene flow and adapted to local climates (e.g. Steane *et al.*, 2015; Jordan *et al.*, 2017; Supple *et al.*, 2018; Ahrens *et al.*, 2019). In one of the first landscape-scale genomic studies in Australia for an understory species, we show that the iconic grass *T. triandra* has very different patterns of connectivity and adaptation compared with its *Eucalyptus* counterparts. Limited dispersal potential and high levels of genetic structuring among remnant populations of *T. triandra* suggests that their adaptability is likely to depend largely on trait plasticity and standing genetic variation that allows for adaptation *in situ*. We provide evidence of genetic and ploidy variation correlated with climate, suggesting that standing genetic variation may be retained within some *T. triandra* populations enabling adaptation to warmer and drier environments emerging under climate change. Indeed, our findings suggest the impacts of climate change may be heterogeneous across the distribution of *T. triandra*. This emphasises the importance of accounting for intraspecific variation, including ploidy, when predicting species responses to new climate challenges. Variability in physiological response to thermal stresses between populations has been established for many plant species (Moran *et al.*, 2016), which may contribute to uneven population responses to thermal stress (Miller *et al.*, 2019). These findings have implications for predicting population responses to climate change, and highlight the importance of interventions (assisted migrations of pre-adapted genotypes) to enhance the resilience of populations showing signs of climate stress given the existence of relatively tolerant populations across the species range.

### Conclusion

Successful establishment of *T. triandra* on three continents from its Asian centre-of-origin is likely due to its ability to swiftly meet the challenges of new environmental conditions through mechanisms unique to the species. Genomic analysis of a species can elucidate broad patterns of structure and provide information about how those patterns are distributed across the landscape. While spatial structure was the major component of the species’ standing genetic diversity, environmental heterogeneity was also a major component driving patterns of diversity, and patterns of neutral genetic diversity have been shown to be affected by natural selection (Phung *et al.*, 2016). Thus, these findings illustrate that the standing genetic variation can provide a basis for adaptation to changing climates and should be incorporated into restoration projects. We were also able to investigate long standing ploidy questions within a landscape genomics context. Notably, we were able to quantify how ploidy might buffer the species from the most severe climate effects in the future. We found that ploidy, along with standing genetic diversity, could be an important part of the puzzle that increases the probability of grassland ecosystem persistence during a period of dramatic change. Our data suggest that we risk underestimating the adaptive capacity of a species if we do not correct for ploidy polymorphisms and we propose that they should be an integral part of management strategies moving forward. Management of multi-ploidy foundational species should focus on a combination of attributes, including genetic variation, intraspecific ploidy polymorphisms, and trait characteristics to develop populations that are resilient to future climate scenarios ensuring ecosystem health, function, and long-term restoration success.

## Data accessibility

Our data will be deposited on Dryad. Including the full SNP data set and population metadata.

## Acknowledgements

We would like to thank Dr Ashley Jones for assistance in the lab for DNA prep for long-read sequences and Dr Robert Godfree for access to a known diploid individual for long-read sequencing. Royal Botanic Gardens Melbourne Friends supported this work through the Helen McLellan Research Grant. JOB and NCA were supported by ARC Centre of Excellence in Plant Energy Biology CE CE140100008.

## Author Contribution

Design of the research was by CA and EJ; collection was performed by CA and EJ along with volunteers; lab work was performed by NA; data analysis was performed by CA; and writing the manuscript was performed by CA, EJ, and AM and all authors contributed to editing the manuscript.

## Supplementary information

**Table S1.** *F_ST_* pairwise table and input for GDM analysis. (tsv file)

**Figure S1.** Elevation of the study area.

**Figure S2.** Histogram of MinION long-read read-lengths and average read quality.

**Figure S3.** Principal components analysis for all 19 bioclim variables.

**Figure S4.** Cross entropy plot to determine the *k*-value for sNMF results.

**Figure S5.** Maps for all four climate variables and ploidy distribution.

