## Supplementary material for "Spatial, climate, and ploidy factors drive genomic diversity and resilience in the widespread grass *Themeda triandra*": Figure S1

Table of contents

| **Table/Figure** | **Description** | **Page #** |
| --- | --- | --- |
| Table S1 | *F*_ST_ pairwise table and input for GDM analysis. | .tsv file |
| Figure S1 | Map of the Great Dividing Range. | 2 |
| Figure S2 | Histogram of MinION long-read read-lengths and average read quality. | 3 |
| Figure S3 | Principal components analysis for all 19 bioclim variables. | 4 |
| Figure S4 | Cross entropy plot to determine *k*-value for sNMF results. | 5 |
| Figure S5 | Maps for all four climate variables and ploidy distribution. | 6 |

Figure S1. Elevation of the study area including the Great Dividing Range, a landscape feature driving temperature and precipitation patterns in eastern Australia. The GDR includes the high elevation areas just inland of the eastern coast. Units are in meters.

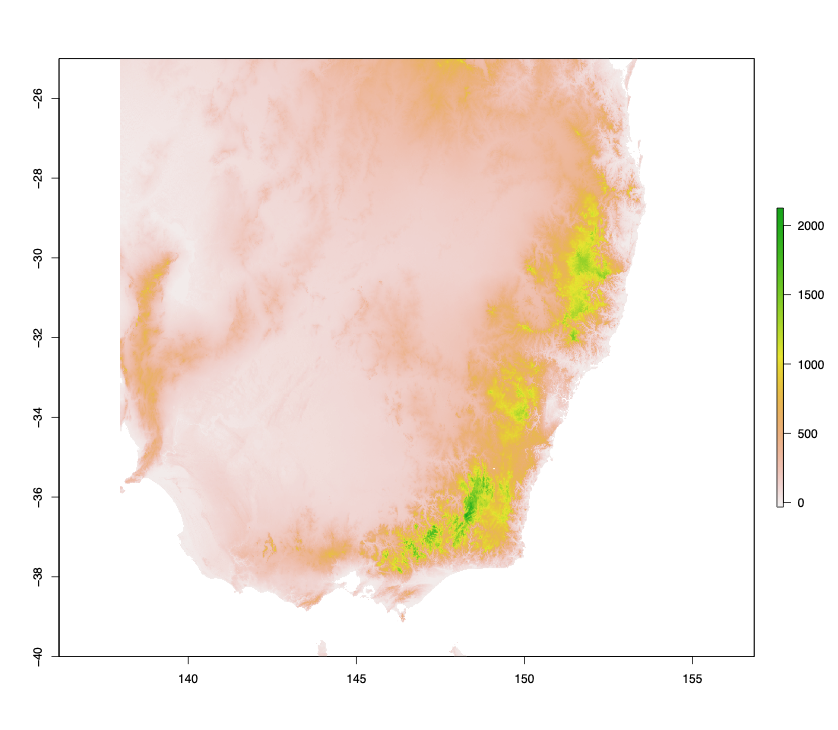

Figure S2. Histogram for the length of sequenced reads from the MinION (top), and the average read quality against read length (bottom).  Reads were up to 140kb long with the majority >30kb.

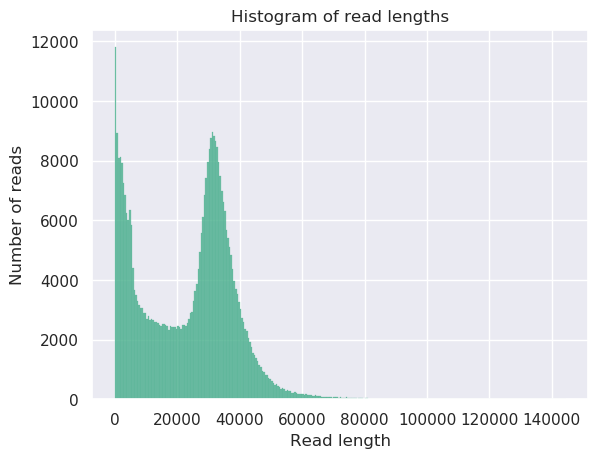

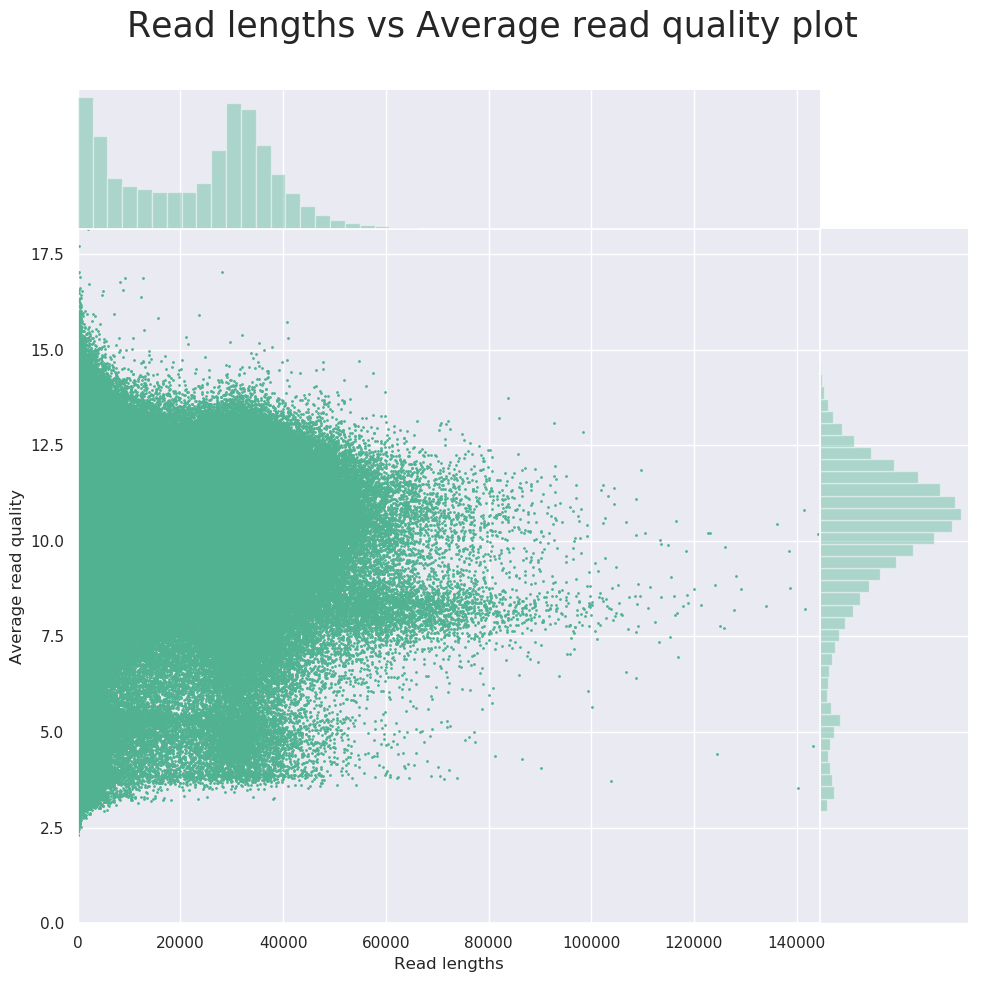

Figure S3. Principal components analysis for all 19 bioclim variables, highlighting correlations between variables. The six climate factors chosen for analyses were temperature mean diurnal range (T_RANGE_; BIO2), maximum temperature of the warmest month (T_MAX_; BIO5), precipitation seasonality (P_SEAS_; BIO15), mean annual temperature (T_MA_; BIO1), mean annual precipitation (P_MA_; BIO12), and precipitation of the driest month (P_DM_; BIO14).

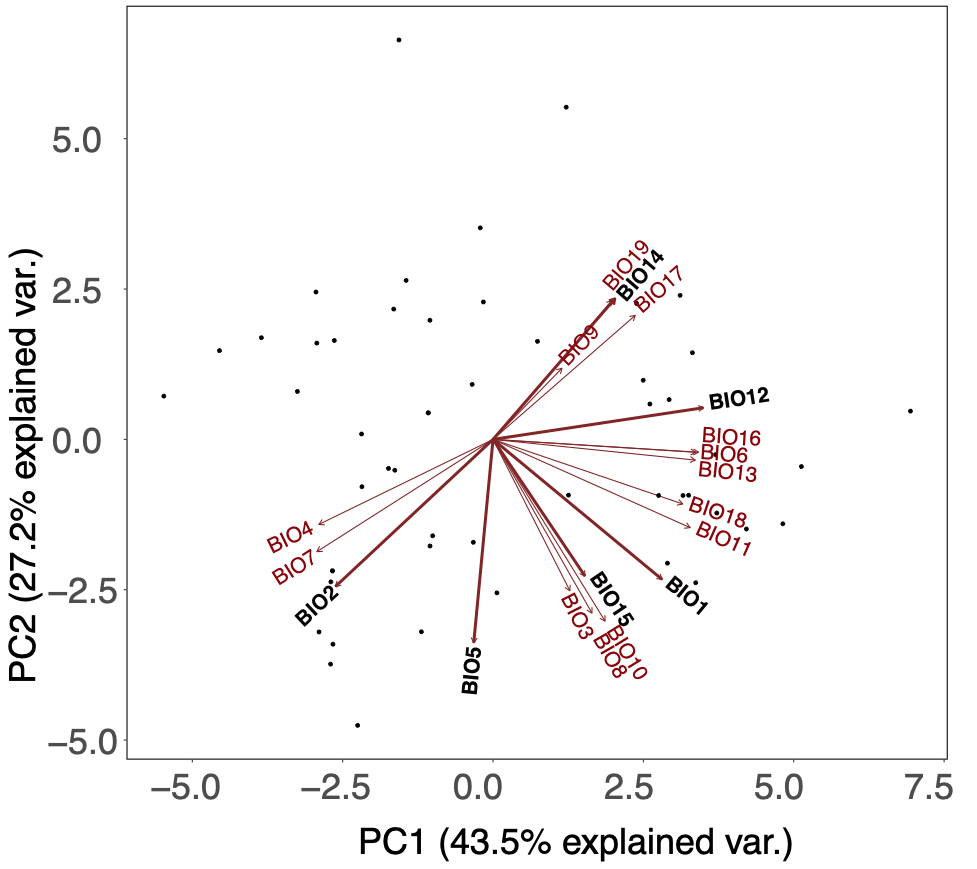

Figure S4. Cross entropy plot for the sNMF results for *k*-values between 1 and 10. A steep decline stops at a *k*-value of 3, therefore we show the sNMF results for *k* = 3.

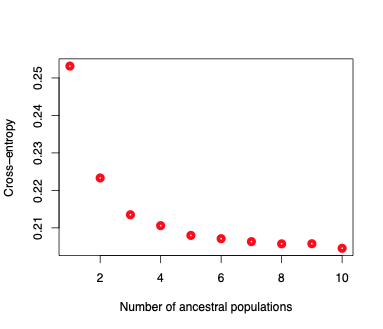

Figure S5. Maps for all four climate variables used in the general dissimilarity visualisation. Climate factors are mean annual temperature (T_MA_; BIO1), maximum temperature of the warmest month (T_MAX_; BIO5), mean annual precipitation (P_MA_; BIO12), and precipitation seasonality (P_SEAS_; BIO15).

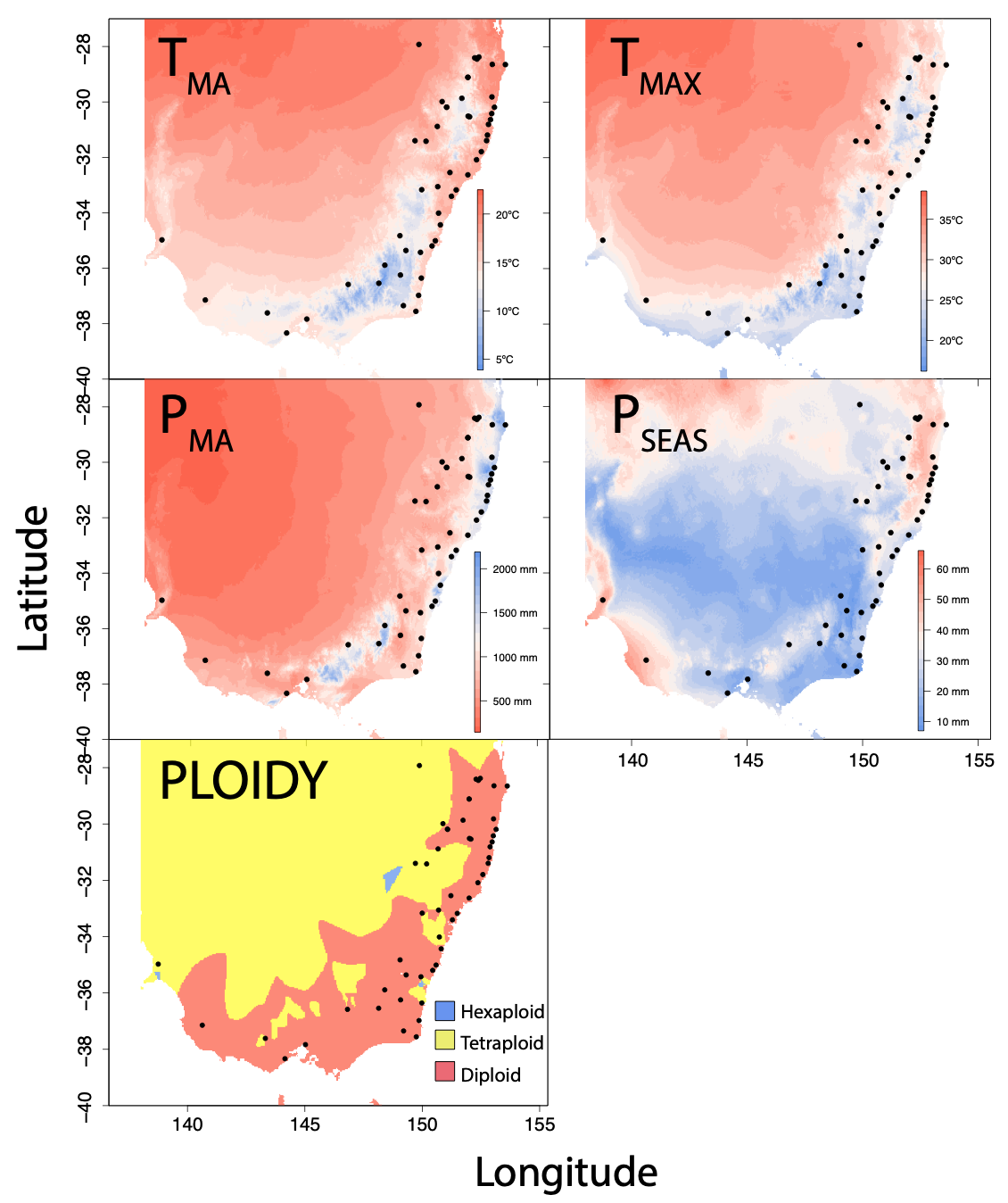
